# Ras protein abundance correlates with Ras isoform mutation patterns in cancer

**DOI:** 10.1101/2021.11.04.467300

**Authors:** Fiona E. Hood, Yasmina M. Sahraoui, Rosalind E. Jenkins, Ian A. Prior

## Abstract

Activating mutations of Ras genes are often observed in cancer. The protein products of the three Ras genes are almost identical. However, for reasons that remain unclear, KRAS is far more frequently mutated than the other Ras isoforms in cancer and RASopathies. We have quantified HRAS, NRAS, KRAS4A and KRAS4B protein abundance across a large panel of cell lines and healthy tissues. We observe consistent patterns of KRAS>NRAS>>HRAS protein expression in cells that correlate with the rank order of Ras mutation frequencies in cancer. Our data provide support for the model of a sweet-spot of Ras dosage mediating isoform-specific contributions to cancer and development. However, they challenge the notion that rare codons mechanistically underpin the predominance of KRAS mutant cancers. Finally, direct measurement of mutant versus wildtype KRAS protein abundance revealed a frequent imbalance that may suggest additional non-gene duplication mechanisms for optimizing oncogenic Ras dosage.

## INTRODUCTION

Ras genes are mutated in ~20% of all human cancer cases ^1^. There are three Ras genes that generate four almost identical proteins: HRAS, NRAS, KRAS4A and KRAS4B ^2^. Despite their similarity, KRAS is far more frequently mutated in cancer. 76% of Ras-mutant cancer patients harbor KRAS mutations versus only 7% with HRAS mutations ^1^. Confirmed explanations for the potent oncogenicity of KRAS have remained elusive since this phenomenon was first noted more than 30 years ago ^3^. How does a family of Ras proteins that share a common set of activators and effectors generate isoform-specific engagement with cancer-associated signaling networks?

At the most fundamental level it must relate to the opportunity and capacity of each Ras isoform to interact with and activate key effector pathways. This is currently best expressed in the Ras “sweet-spot” model that suggests that Ras dosage will be a major factor in influencing the availability of individual Ras family members to engage cancer pathways ^4^. This model is an iteration of wider “Goldilocks” models describing oncogenic dosing contributions to cancer ^5^. It suggests that there is an optimal level of Ras expression in each tissue and genetic context that will promote cancer. Lower levels of expression will be insufficient to initiate tumorigenesis, whilst too much Ras will induce oncogenic stress. Consistent with this, high levels of Ras or downstream Raf-MAPK activation are known to induce senescence and cell death ^6, 7, 8, 9^

Ras dosage is known to be important for KRAS-mediated progression of pancreatic and breast cancer and KRAS and NRAS contributions to myeloid malignancies ^9, 10, 11, 12^. The mechanistic basis linking Ras isoforms, Ras dosage and cancer mutation patterns was potentially provided by the observation that the KRAS gene is enriched in rare codons ^13^. Rare codons limit protein translation efficiency ^14, 15^ and optimizing codons in the KRAS gene locus did indeed result in higher KRAS protein expression ^13^. Consistent with the sweet-spot model, higher KRAS expression reduced carcinogen-induced tumorigenesis in mice and altered engagement with cancer signaling pathways ^13, 16^, ^17, 18^. As a result, it was suggested that KRAS expression is optimal in most contexts, whereas HRAS and NRAS expression is too high ^4, 13^.

Importantly, HRAS, KRAS and NRAS protein abundance was never formally measured in these studies to see whether they conformed with the predicted influence of rare codons. Moreover, rather than exhibiting limited expression, KRAS is actually far more frequently amplified in tumors than the other isoforms ^19^. KRAS mRNA represents 70-99% of all Ras transcripts in mouse tissues ^20^, and quantitation of Ras protein abundance identified KRAS to be the most abundant isoform in a selection of cancer cell lines ^21, 22^. Therefore, whilst the evidence for Ras dosage influencing tumorigenesis is compelling, the proposed rare codon link between KRAS and cancer mutation patterns remains contentious.

In order to address this, we have quantified Ras protein abundance across a large panel of cell lines and healthy tissues. Whilst our insights do not agree with the predicted influence of rare codons on KRAS protein expression versus other isoforms, we do observe consistent patterns of Ras isoform expression suggesting that relative dosage is an important feature of their biology and disease association. We also observe an imbalance in the abundance of proteins expressed from mutant versus wild type alleles in some cell lines that suggests the existence of additional mechanisms for achieving disease-associated amplification. These datasets and the patterns observed have broad applications in experimental design, network analysis and understanding the contributions of Ras isoforms to normal and disease-associated biology.

## RESULTS

The patterns of Ras mutations in cancer are suggested to be influenced by rare codon-mediated differences in protein expression. This was established in mice where the KRAS gene is enriched in rare codons versus HRAS ^13^. The codon adaptation index (CAI) is a measure of synonymous codon usage bias in a DNA sequence ^23^. CAI analysis of human Ras exons reveals that KRAS is not an outlier like in mice, instead it exhibits a similar enrichment of rare codons as NRAS (Figure 1A). This suggests that rare codon-mediated limitation of Ras protein expression is not responsible for the much higher representation of mutants of KRAS than NRAS in human cancer patients.

**Figure 1.**
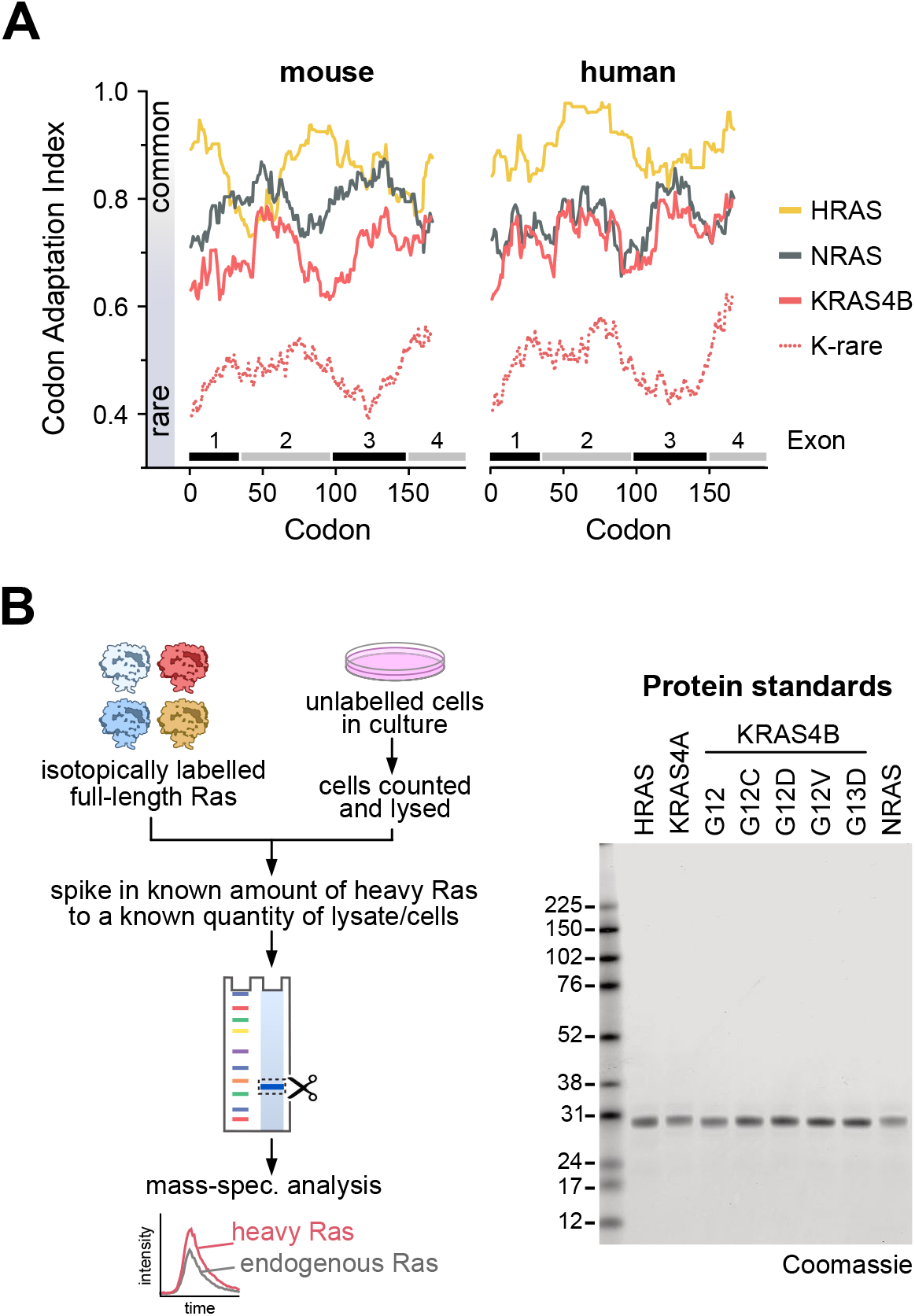
Ras codon bias and methods for protein quantitation. Rare codons are equally enriched in NRAS and KRAS in humans (**A**). Schematic for absolute quantitation of Ras protein abundance (**B**). Coomassie blue staining of 200 ng of isotope labelled “heavy” Ras protein standards indicates high purity suitable for precise quantitation and ratiometric comparison with endogenous Ras proteins.

To formally measure Ras dosage, we employed a mass-spectrometry based protein standard absolute quantitation (PSAQ) approach to determine Ras isoform protein copy number per cell (Figure 1B) ^22, 24, 25^. High purity isotope-labelled, fulllength Ras standards are quantified and known amounts spiked into cell lysates derived from a known number of cells. Spike-in at this early stage improves accuracy by ensuring normalization of potential variables associated with subsequent sample processing. Ras isoform pre-enrichment steps are not required, this removes a major source of potential error associated with non-quantifiable differences in immunoprecipitation efficiencies. Following fractionation and trypsin digestion, diagnostic peptides for each Ras isoform together with a pan-Ras peptide shared by all isoforms (Supplementary Figures 1 & 2A) are detected by mass spectrometry and quantified. We observe clear cross-correlation of Pan with H+N+KA+KB peptides across all wild type cell lines indicating that all peptides are quantitative with a high degree of accuracy (Supplementary Figure 2B). The pan-Ras peptide includes codon 12 and 13; therefore, it will only report total Ras in wild type cells, and only wild type Ras abundance in mutant Ras cell lines. To allow precise quantitation of mutant Ras abundance, relevant standards of four common Ras mutants were also prepared (Figure 1B). These standards are linearly responsive over dynamic ranges relevant to the range of Ras concentrations observed in cell lines (Supplementary Figure 1). All proteotypic Ras peptides were quantified using a minimum of three transitions (Supplementary Figure 1). Ratiometric comparison of peptides from the mass-shifted isotope labelled standard versus the endogenous protein enable accurate determination of lysate protein abundance that can be integrated with cell counts to calculate protein copy number per cell.

PSAQ was applied to 64 commonly used mouse and human cell lines. All data are derived from three independently processed cell samples where a common Ras standard was spiked into all samples to allow direct comparison between all cell lines. Total Ras abundance derived from H+N+KA+KB peptide measurements ranges over an order of magnitude from ~50,000-600,000 proteins per cell (Figure 2, Supplementary Table 1). This highly correlates with a proxy for cell size (Supplementary Figure 2C), and we observe that total Ras abundance occupies a relatively narrow -2-3-fold range centered on the linear correlation trend line. Both mouse and human cell lines exhibit similar levels of Ras abundance.

**Figure 2.**
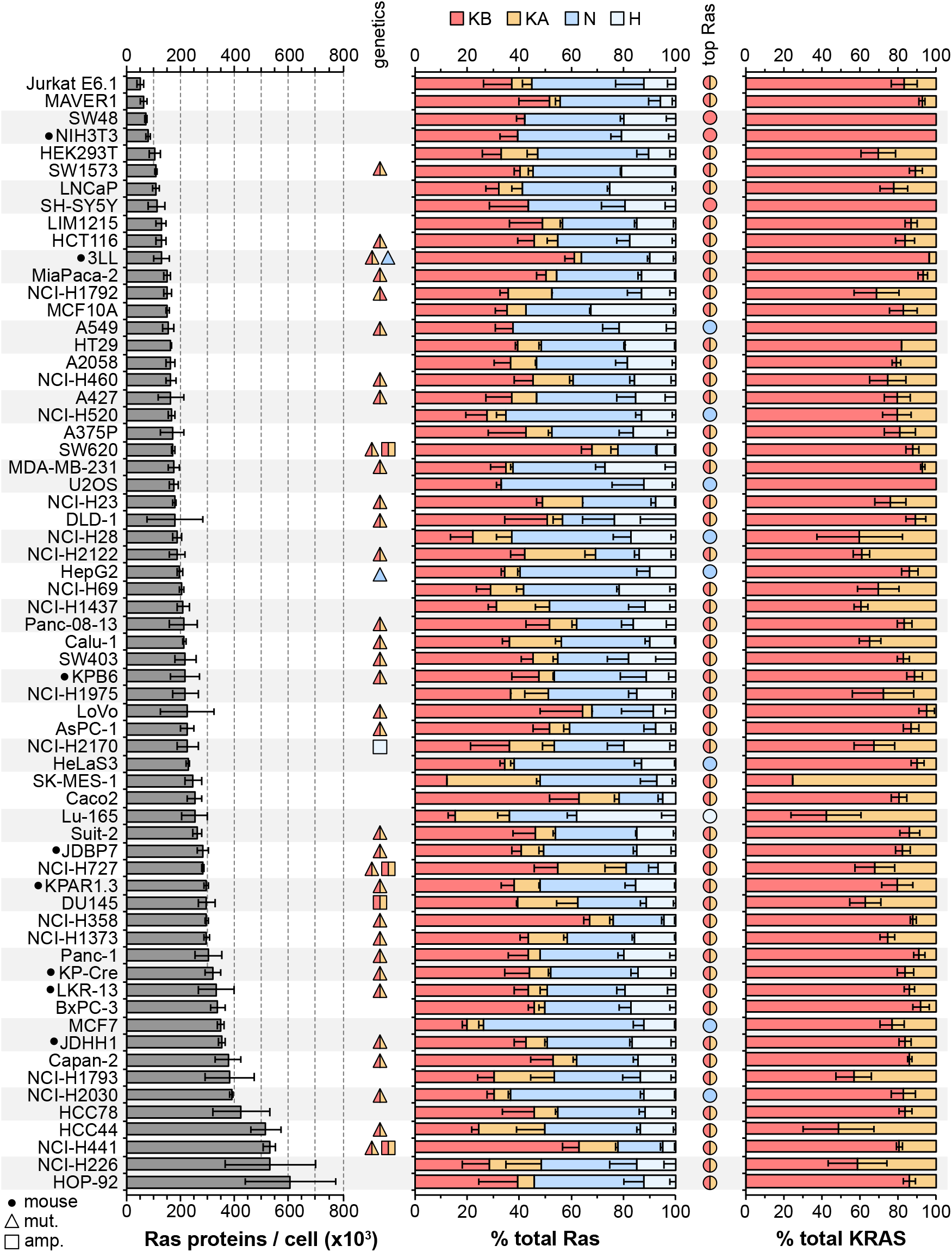
Ras protein abundance in a panel of cell lines. Ras proteins are highly abundant, the significant variation in total Ras abundance correlates with cell size (Supplementary Figure 2). Aggregate KRAS abundance averages ~50% across the cell panel, in most cell lines KRAS4B expression exceeds the other isoforms. Measurements represent mean ± SEM of n=3 independently processed and analyzed cell samples unless otherwise indicated in Supplementary Table 1.

Quantitation of Ras isoform protein abundance reveals a relative rank order of KRAS4B>NRAS>>HRAS>KRAS4A. KRAS4B is the dominant Ras isoform in 44/64 cell lines, whilst KRAS gene products totaled together represented the most abundant Ras in 51/64 of cell lines (Figure 2). The upper limit of KRAS contribution to total Ras was ~80% (average 52%), for NRAS it was ~60% (average 33%), and for HRAS it was ~40% (average 15%). Whilst data on whether there is Ras amplification in these cell lines are not comprehensive, all examples of known KRAS amplification correlated with PSAQ measurement of very high relative percentages of KRAS in these cell lines.

KRAS4A averages only ~17% of total KRAS abundance (Figure 2, Supplementary Table 1). This isoform was the most challenging to detect, it was completely undetectable in five cell lines and wasn’t detected in at least one replicate of a further twenty-four. We think that this is due to its abundance being low and close to the sensitivity limit of our assay rather than evidence of systematic under-estimation of KRAS4A abundance. Our confidence in the accuracy of our measurements is supported by observations in wild type cell lines that total Ras abundance calculated from the pan-Ras peptide closely correlates with total Ras calculated from summing the isoform-specific peptides (Supplementary Figure 2).

The consistent patterns of Ras isoform abundance across a large panel of normal and cancer cell lines derived from two species are compelling. However, it is formally possible that Ras levels may have been influenced by disease of origin, cell derivation and/or culture conditions. To address this, we profiled a panel of tissues freshly derived from three healthy adult CD1 mice. We observed a 4-fold range in total Ras abundance across the tissues (Figure 3A). Our previous studies quantified Ras isoform transcript abundance in the same mouse strain using RT-PCR ^20^. Our measurements of total Ras protein abundance broadly correlated with these measures of transcript abundance, with brain and lung again displaying the highest levels of Ras (Figure 3B). Similar to the cell line observations, KRAS is the most abundant Ras in every mouse tissue profiled (Figure 3A). In all tissues except skeletal muscle KRAS is more abundant that HRAS and NRAS combined. However, in contrast to what was observed in the cell lines, HRAS was more abundant in mouse tissues than NRAS.

**Figure 3.**
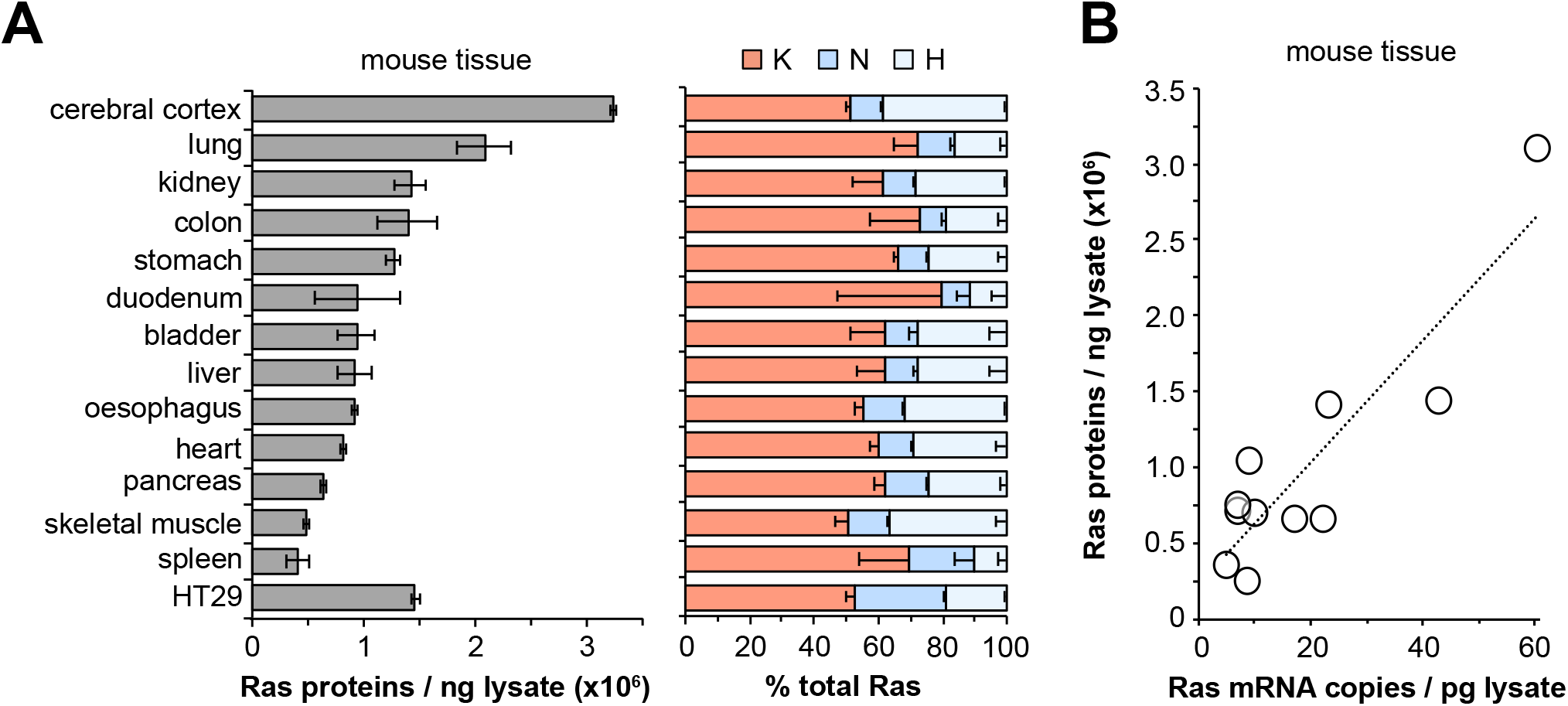
Ras protein abundance in tissues. Total Ras (pan-Ras peptide) abundance varies 2-3-fold across mouse tissues (**A**). KRAS is the most abundant isoform in all tissues. Total Ras protein and transcript abundance correlate in mouse tissues (**B**). All protein measurements represent mean ± SEM of tissues derived from n=3 adult mice. Transcript data derived from ^20^.

Many of the cell lines that we profiled harbored KRAS mutations (Supplementary Table 1). Previous work from our lab suggested that there might be a predominance of mutant versus wild type protein abundance in some members of a panel of isogenic SW48 cells ^22^. To investigate this phenomenon, we generated high purity isotope-labelled Ras standards for four of the most common KRAS mutants (Figure 1). Peptides derived from amino acids 6-16 normally used for Pan-Ras quantitation now contain the mutation, allowing specific ratiometric measurement of mutant Ras protein copy numbers.

A selection of cell lines with relevant heterozygous KRAS mutations were profiled for mutant protein abundance and this was compared with KRAS4A + KRAS4B abundance to determine relative mutant versus wild type percentages (Figure 4). For G12D, G12V and G13D mutant cell lines we observed at least one example of a cell line with approximately equivalent proportions of mutant versus wild type. However, in every case we also observed imbalanced frequencies favoring higher mutant Ras protein abundance. For G12C mutant cell lines this was even more apparent with every cell line tested exhibiting mutant predominance. Together, these data suggest that mutant Ras signaling somehow differentially regulates mutant versus wild type KRAS protein dosage.

**Figure 4.**
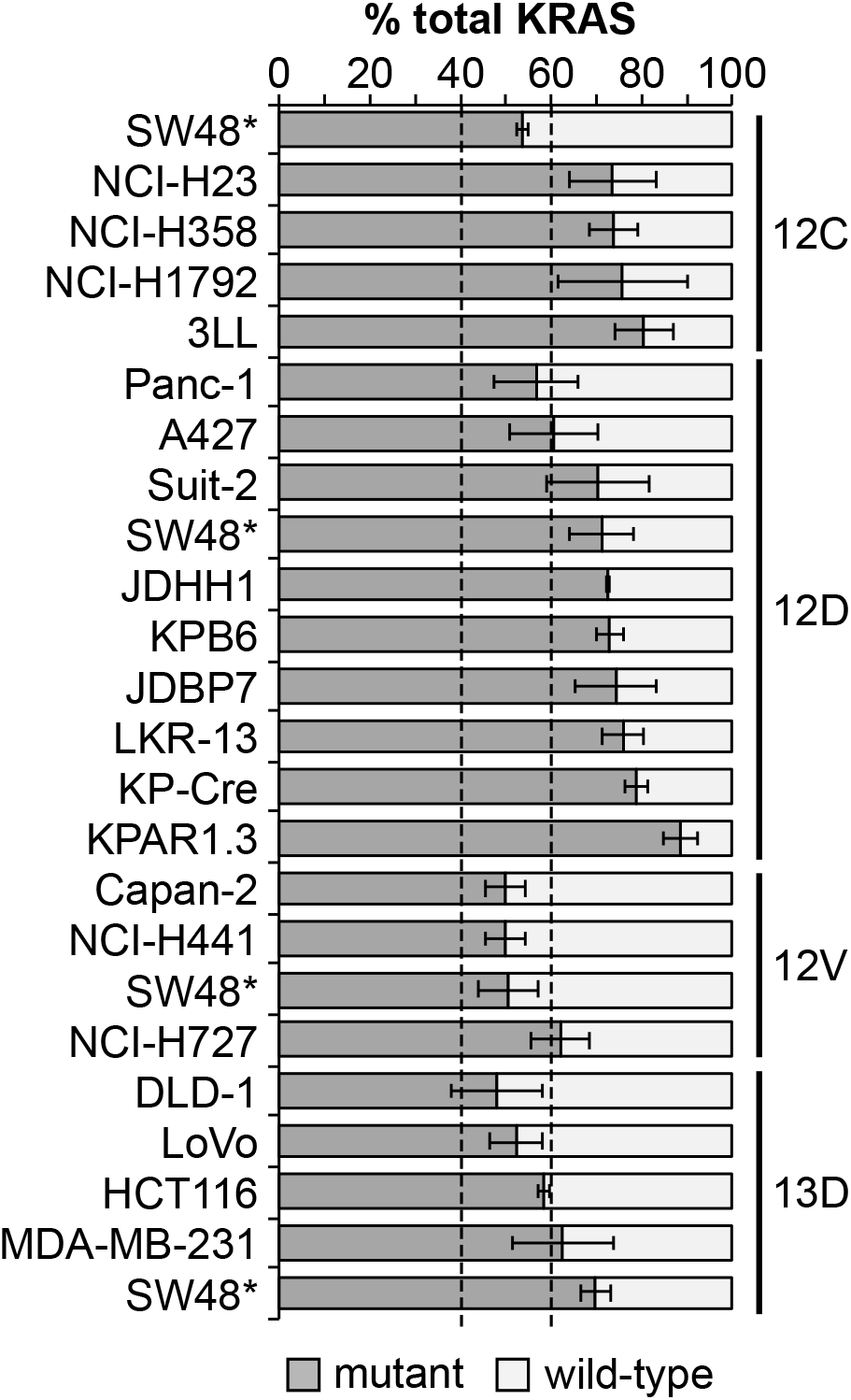
Imbalanced ratios of mutant : wild type Ras proteins. Direct quantitation of mutant versus wild type KRAS proteins reveals a frequent excess of the mutant protein in heterozygously mutated cell lines. All measurements represent mean ± SEM of n=3 independently processed and analyzed cell samples.

## DISCUSSION

We have quantified Ras protein abundance in a large panel of cell lines and healthy tissues to help understand relative dosage contributions of Ras isoforms to cancer and development. The method that we have employed overcomes major issues that hinder accurate protein quantification because it does not rely on pre-enrichment steps and the Ras standards experience the full sample processing pipeline. Our measurements reveal a wide range of total Ras abundance (50,000-600,000 copies per cell). When normalized to cell size this narrowed to a 2-3-fold difference in the highest versus lowest values of total Ras abundance for a given cell size. Total Ras abundance for an averaged sized cell was ~200,000 copies per cell. This would place Ras in the top 20% of all proteins based on global estimates of mammalian cell proteome copy number ^26, 27, 28^. It also means that Ras is a relatively abundant node within the wider Ras signalling network ^28^. Copy number per cell is a relatable measure of abundance; however, actual Ras concentrations will be influenced by relative partitioning between membrane, cytosol and subcellular compartments. Therefore, a more refined understanding of biologically relevant Ras isoform dosage will also need more accurate quantification of Ras localization.

Across cell lines and normal tissues, we observed a consistent pattern of KRAS being the most abundant isoform. This corroborates and significantly extends previous studies that also identified KRAS to be the most abundant Ras isoform in a small panel of cell lines ^21, 22, 29^. Our method was also capable of quantifying KRAS splice variants. The most comprehensive previous analysis of the expression of these isoforms applied QPCR to 30 human cell lines and 20 colorectal cancer samples ^30^. It found that KRAS4A represented 5-50% of total KRAS transcripts. Protein abundance was quantified using Western blotting in only one cell line, and in contrast to their transcript data, KRAS4A appeared to be at least twice as abundant as KRAS4B in our measurements in HT29 cells ^30^. In our study we directly measured KRAS isoform protein abundance in 64 cell lines representing a diverse range of tissues. KRAS4A was clearly the minor isoform averaging ~17% of total KRAS. In HT29 cells we found KRAS4A to be at the limits of detection and estimate that it represents only ~10% of total KRAS.

Intriguingly, the rank order of Ras isoform abundance differed between cell lines and mouse tissues. In all mouse tissues except spleen HRAS was more abundant than NRAS. In contrast, in all cell lines apart from MCF10a, DLD1 and LU165 cells NRAS was more abundant than HRAS. The same batches of Ras standards were used for cell and tissue analysis. HT29 cells were included in all PSAQ sample runs and retained the consistent pattern of NRAS>HRAS seen in all previous runs; therefore, we are confident that the mouse tissue data represent a true reflection of Ras abundance in these samples. The switch in abundance may represent general species differences although we note that our cell panel included eight mouse lines that all displayed the same trends as their human counterparts. Alternatively, it may represent adaptive changes between normal versus the disease associated states that our cells were derived from. Further investigation of normal human tissues and relevant mouse models before and after disease initiation are required to test this.

The patterns of Ras isoform expression that we observe potentially inform our understanding of why KRAS is more frequently mutated in cancer. Our results challenge the central tenet of the rare codon model that predicted that KRAS expression would be limited relative to the other Ras isoforms ^4, 13, 16^. We found that in humans, KRAS and NRAS share similar compositions of rare codons; whilst in cells and tissues, KRAS was clearly the most abundant isoform. Although this means that rare codons are not the mechanistic basis underpinning observed differences in Ras protein dosage, we note that KRAS mRNA levels are disproportionately higher than their relative protein abundance ^20^. This may be a compensatory mechanism for overcoming reduced translational efficiency. KRAS transcripts also include extensive untranslated regions compared to the other Ras isoforms. It is tempting to speculate that the abundance of rare codons in KRAS might be a mechanism for achieving high transcript copy numbers that could be biologically meaningful via genetic rather than protein-based mechanisms. Rare codons may also provide a temporal control over Ras expression since they are enriched in genes exhibiting increased expression during proliferation ^31, 32^.

Whilst our data mean that we do not now think that rare codons explain why KRAS is more often mutated in cancer, the rare codon experiments and work by others have conclusively demonstrated that Ras dosage has a significant influence on Ras oncogenic potential ^9, 10, 11, 12, 16^. The Ras sweet-spot model remains the most compelling theory for explaining Ras mutation patterns ^4^; however, our cell and tissue data suggest that HRAS and NRAS expression levels in most contexts are likely to be insufficient rather than oncogenically stressful. Our revised view is summarized in Figure 5. We suggest that KRAS is the most frequently mutated Ras isoform in cancer because it has a higher level of expression that is closer to the sweet-spot compared to the other isoforms. This is also consistent with the highly biased pattern of KRAS amplification in cancer compared to the other isoforms ^19^, suggesting higher levels of KRAS are more often hitting the sweet-spot, whereas amplification of the other isoforms remains insufficient in most contexts. Notably, KRAS mutation frequency is disproportionately higher than NRAS compared to their relative abundance. This suggests that there is a step change in oncogenic potential in the 30-50% zone of total Ras abundance where these isoforms overlap.

**Figure 5.**
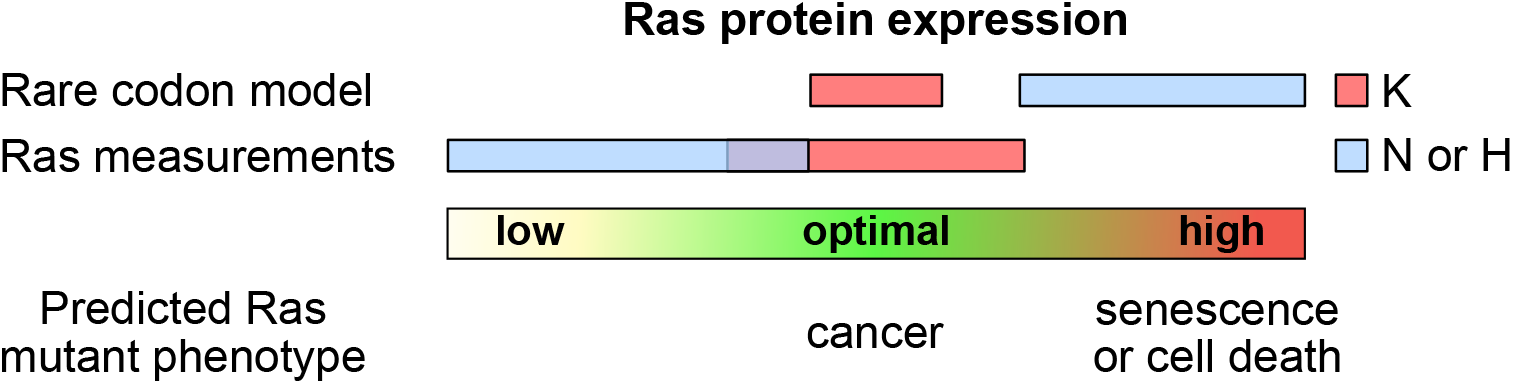
A revised view of the Ras sweet-spot model. Our data suggest that protein abundance of HRAS and NRAS is too low rather than too high for promoting oncogenesis in most contexts.

Ras dosage is likely to be more nuanced than measurements of total abundance and it will be interesting to understand if total versus compartment-specific Ras are the critical determinants of the Ras sweet-spot. The type and amount of mutant Ras are also likely to be influential. Mass spectrometry currently represents the only method for directly measuring mutant Ras protein abundance ^33^. Our analysis repeatedly observed a preponderance of mutant versus wild type KRAS protein in heterozygous cell lines. Confidence in the accuracy of mutant Ras quantitation is increased by corroborating data from other studies. PROTAC and siRNA mediated loss of G12C mutant KRAS in NCI-H23 and NCI-H1792 cells resulted in loss of >75% of total KRAS, analogous to our estimates of high ratios of mutant KRAS in these cell lines ^34, 35^.

Our mutant Ras data provide evidence of fine tuning of cancer-relevant Ras dosage. The dynamic modulation of optimal total and mutant Ras dosage has been elegantly demonstrated in the characterization of allelic imbalances in mutant versus wild type KRAS during tumor outgrowth and in response to therapy in a mouse model ^12^. The increased ratio of mutant versus wild type KRAS abundance that drove the cancer responses in that study is analogous to what we also observed. Our data suggest an alternative mechanism to gene duplication that could further help to titrate mutant Ras contributions to tumorigenesis.

Ras dosage is also likely to explain Ras isoform-specific contributions to development. KRAS4B is the only essential isoform, deletion results in embryonic lethality as a result of heart defects and cardiovascular problems ^36, 37, 38^. Gene swap experiments revealed that HRAS could compensate when driven from the KRAS locus ^39^, implying that it was expression rather than unique signaling abilities of Ras isoforms that was important. Our data reveal that KRAS represents 50-75% of total Ras across a range of mouse tissues including the heart. Therefore, even in the double HRAS/NRAS knockout mice that were able to generate viable offspring ^40, 41^, all tissues would still contain ≥50% of normal Ras levels.

In summary, we have generated a comprehensive atlas of Ras isoform protein abundance in tissues and commonly used cell lines. Accurately quantifying Ras protein abundance will help parameterize models of signaling networks and inform cell-based Ras studies. Our data reveal new features of Ras biology and challenge and refine models explaining the pattern of Ras mutations in cancer and isoform-specific Ras contributions to development.

## MATERIALS AND METHODS

### Cell lines, counting and lysis

Verified cell lines were obtained from sources indicated in Supplementary Table 1 Prior to use, all cells tested negative for mycoplasma using an e-Myco Plus kit (Intron Biotechnology. Cells were grown to 60-100% confluence and harvested by trypsinisation and counted using a Countess II FL automated counter (Thermo). Cell lines with large cells were manually counted with a haemocytometer. Pellets corresponding to 1.5-70 x 10^6^ cells were washed twice with ice cold PBS by centrifugation, snap frozen using liquid nitrogen and stored at −80 °C. Pellets were thawed on ice and lysed in NP40 lysis buffer (50 mM Tris, pH 7.5, 150 mM NaCl, 1% NP40 substitute, 1/250 mammalian protease inhibitor cocktail (Sigma)). Lysate protein concentration was determined using Pierce BCA assay. n=3 independently prepared cell pellets were used for PSAQ analysis.

### Mouse tissue lysates

Mouse tissues were harvested from adult male B6/129S mice, snap frozen in liquid nitrogen and stored at −80 °C. Tissue pieces of ~30-50 mg were crushed with a mini pestle chilled on dry ice before lysis in RIPA buffer (50 mM Tris, pH7.5; 150 mM NaCl; 1 x Triton x100; 0.1% SDS, 1% sodium deoxycholate, 1/250 mammalian protease inhibitor cocktail (Sigma)). Samples were sonicated on ice and lysate cleared by centrifugation at 17,000 x g. Protein concentration was determined using Pierce bicinchoninic acid (BCA) assay. Tissues from n=3 adult mice were used for PSAQ analysis.

### Production of recombinant heavy labelled His-Ras proteins

His-Ras vectors (^22^ or kindly supplied by Dominic Esposito, NCI Ras Initiative) were transformed into AT713 bacteria (Yale E.Coli Genetic Stock Centre) that are auxotrophic for Lysine, Arginine and Cysteine. Heavy (L-lysine-U-^13^C_6_-^15^N_2_ [+8Da]), L-arginine-U-^13^C_6_-^15^N_4_ [+10Da])-labelled His-Ras proteins were prepared exactly as described ^42^. Ras proteins were quantified using the BCA assay. Mass-spectrometry was used to confirm full isotopic labelling efficiency and to verify the accuracy of quantification of the protein concentration of each Ras variant by pairwise comparison with a known quantity of unlabelled His-KRas4B protein using the shared Pan-Ras peptide. A single master stock each for wild type and mutant heavy Ras standards was prepared for use in PSAQ analysis of all relevant cell and tissue samples. Proteins were boiled in sample buffer (60 mM Tris–HCl (pH 6.8), 2% SDS, 5%β-mercaptoethanol, 0.02% bromophenol blue, 9% glycerol), aliquoted and stored at −80 °C. Final spike-in amounts of wild type Ras were 2 ng His-KRas4B, 1 ng His-KRas4A, 1 ng His-HRas and 1 ng His-NRas for cell analysis; in tissue this was supplemented by an additional 2 ng of His-KRas4A. Quantitation of mutant Ras used spike-ins of 2 ng each of wild type and mutant His-KRas4B.

### Preparation of PSAQ samples

20 μg of cell lysate or 40 μg of mouse lysate containing spike-ins of the relevant heavy Ras standard were fractionated using SDS PAGE. The region of the gel containing endogenous and His-Ras standards was excised and dissected into ~1 mm gel cubes. These were reduced using 10 mM DTT in 100 mM Ammonium bicarbonate (Ambic) for 1 hour at 56 °C, alkylated in 55 mM iodoacetamide for 30 mins at room temperature, quenched with 10 mM DTT for 5 minutes at room temperature, then digested with 5 ng / μl Trypsin Gold (Promega) in 9% Acetonitrile and 40 mM Ambic, overnight at 37 °C. Trypsin was quenched using Formic acid and extracted peptides dried using a SpeedVac. Peptides were subjected to C18 desalting using an Agilent 1260 Infinity LC system equipped with an MRP-C18 Hi-recovery column (Agilent, USA) before SpeedVac drying.

### Mass spectrometric quantification of Ras isoforms

Desalted samples reconstituted in 0.1% formic acid were delivered into a QTRAP 6500 (Sciex) *via* a Dionex U3000 nano-LC system (Thermo) mounted with a NanoAcquity 5 μm, 180 μm x 20 mm C_18_ trap and 1.7 μm, 75 μm X 100 mm analytical column (Waters) maintained at 40 °C. A gradient of 2-50% acetonitrile / 0.1% formic acid (v/v) over 45 minutes was applied to the columns at a flow rate of 300 nL / minute. The NanoSpray III source of the mass spectrometer was fitted with a 10 μm inner diameter PicoTip emitter (New Objective). The mass spectrometer was operated in positive ion mode using Analyst TF1.6 software (Sciex) and the MIDAS approach (MRM-initiated detection and sequencing) was used to quantify and confirm the identity of the analytes of interest. The optimized transitions are shown in Supplementary Table 1; dwell time for each transition was 20 ms. The charge status of each precursor ion was determined using an enhanced resolution scan at 250 Da/s and up to 3 MS/MS scans at 10,000 Da/s were triggered with dynamic fill time. This gave a total cycle time of 3.9 s.

### Analysis of MRM

Area under curve was extracted for at least 3 combined transitions in Analyst software (Sciex), and further analysis performed in Excel. Briefly, ratios of Light:Heavy for each peptide were used to determine the quantity of peptide present per lane in moles, before conversion into copies per cell using the number of cells counted per μg lysate produced. Three repeats of each experiment were performed, and for each condition a mean was calculated from at least 2 values for inclusion in the final data. For KRas4B up to 6 values were obtained as this was analysed in both “wildtype” and “mutant” heavy spike ins. The same analysis was performed for mouse tissue lysates, although copies per ng total protein were calculated. For mouse tissues, KRas4A and 4B proved to be less consistent than NRas and HRas, so a combined value for KRas copies per ng total protein was calculated by subtracting copies of NRas and HRas from the pan-Ras copy number.

## ACKNOWLEDGEMENTS

This work was supported by funding from NWCR (CR1166) and the Wellcome Trust (WT203983).

## DATA AVAILABILITY STATEMENT

The authors declare that all data that support the findings of this study are available within the paper and supplementary files. Raw mass-spectrometry data are available from PASSEL: http://www.peptideatlas.org/PASS/PASS01706

---

**Supplementary Figure 1.**
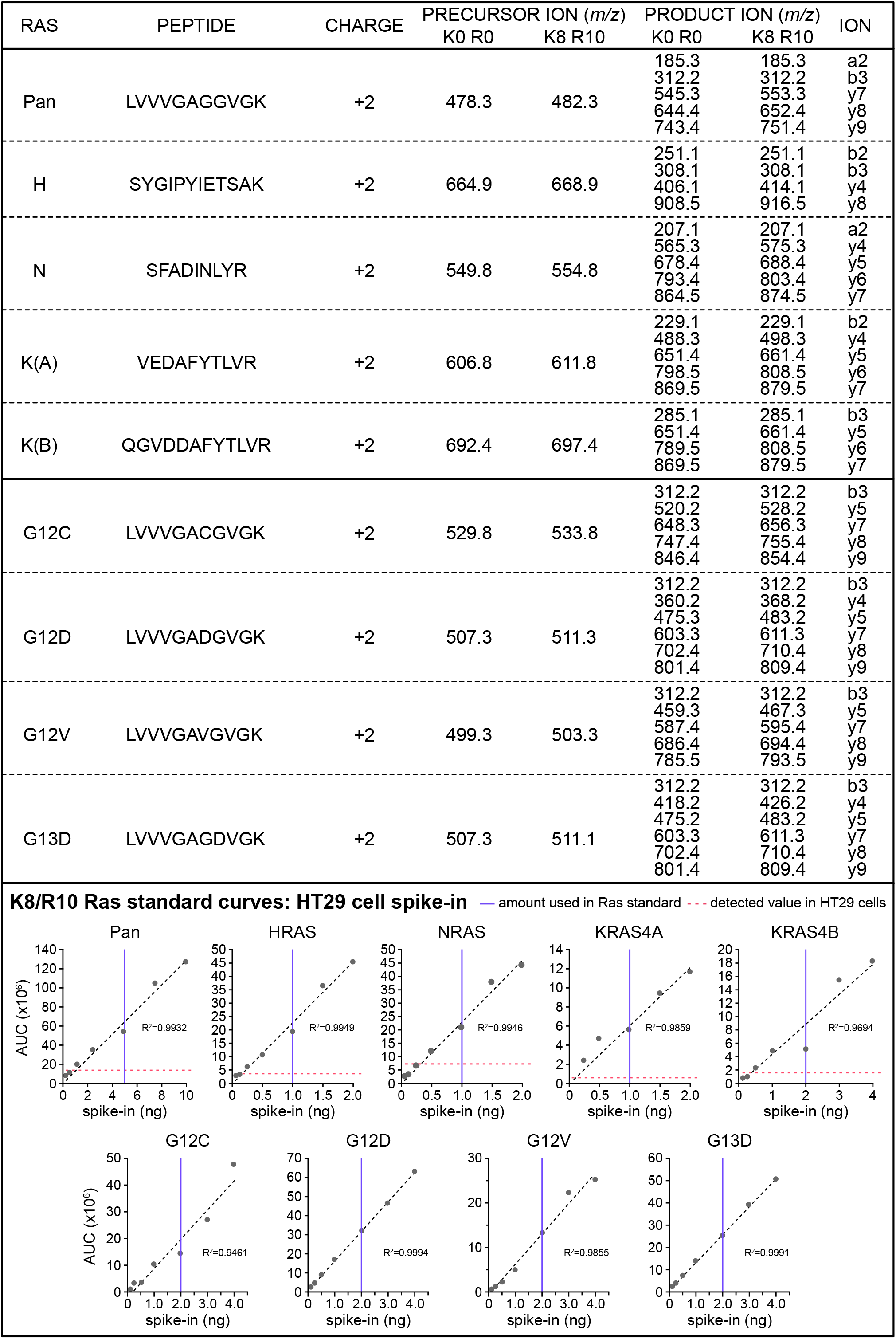
Transitions for proteotypic Ras peptides and integrated detection sensitivity. Transitions chosen for endogenous (Lys0 Arg0) and isotope-labelled (Lys8 Arg10) Ras peptides. A linear relationship is observed (R^2^ > 0.94) for detecting and quantifying Ras isoforms over a wide range corresponding to those observed for endogenous Ras in cell lines.

**Supplementary Figure 2.**
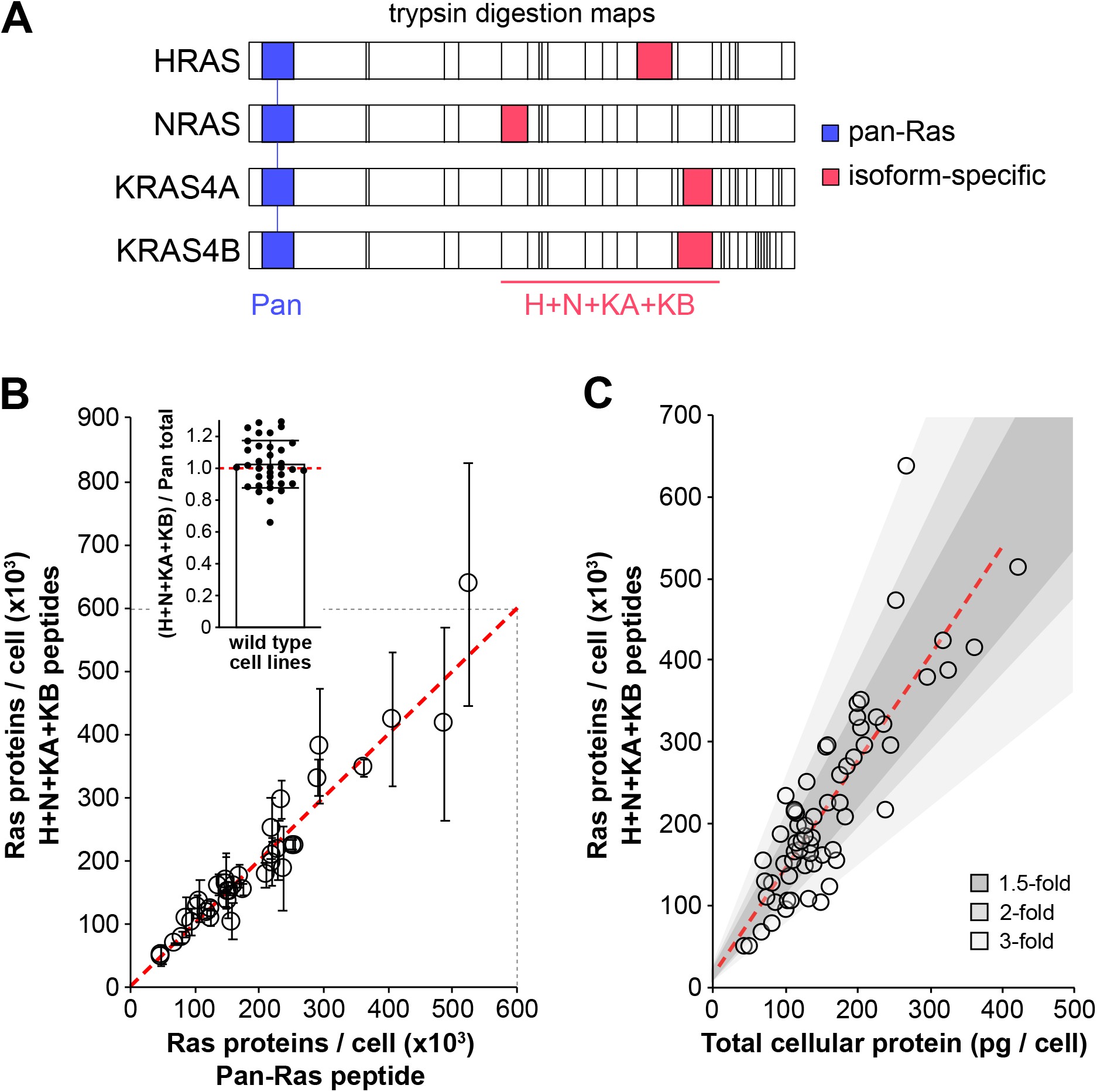
Correlating Ras peptides and total Ras with cell size. Tryptic digestion maps of Ras isoforms with the locations of pan and isoform-specific peptides indicated (**A**). The pan-Ras peptide measure total Ras in wild type cells. Pan and H+N+KA+KB peptides closely correlate in all wild type cell lines; their ratios average 1.00 as expected if quantification of all peptides is accurate (**B**). Average protein abundance per cell is used as a proxy for cell size. Total Ras abundance and cell size highly correlate (R^2^=0.73) (**C**). Almost all cell values lie within a 3-fold range centered on the trend line. All measurements represent mean ± SEM of n=3 independently processed and analyzed cell samples.

**Supplementary Table 1.** *Ras protein measurements in cells and tissues*. Means ± SEM of n=3 independently processed and analyzed samples from 64 cell lines and 13 tissues. All source data are included.

